# Geographical restructuring of *Arabidopsis thaliana*’s genetic makeup in the Iberian Peninsula due to climate change based on genetic cluster membership

**DOI:** 10.1101/091686

**Authors:** Arnald Marcer, Marie-Josée Fortin, F. Xavier Picó

## Abstract

**Aim:** To assess the effects of climate change on genetic lineages of *Arabidopsis thaliana* at the *admixed* population level by directly modelling genetic cluster membership values to predict potential genetic cluster memberships across the Iberian Peninsula.

**Location:** Iberian Peninsula

**Methods:** We used a dataset of 274 accessions structured in four genetic clusters as inferred from 250 nuclear single-nucleotide polymorphisms with Bayesian clustering methods. We predicted the change in percentages of genetic cluster membership at a population level and the changes in potential suitability across the study area by combining parametric (Beta regression) and non-parametric (Recursive trees) methods.

**Results:** Climate change will affect genetic lineages of *Arabidopsis thaliana* differently. Genetic clusters GC1 and GC2 will suffer a substantive reduction of their respective suitable areas while GC3 and GC4 will expand northward. At the population level, except for GC4, the rest of the lineages will undergo a genetic turnover for many of their populations.

**Main conclusions:** *A. thaliana* in the Iberian Peninsula will undergo a major internal genetic restructuring and range change due to climate change. Genetic lineages of *Arabidopsis thaliana* in the Iberian Peninsula will be affected differently which reinforce the need for taking into account intraspecific genetic variation when modelling species distribution. Despite limited predictive power of individual statistical models, the combination of distinct models can compensate this shortcoming.

## INTRODUCTION

The intraspecific response of species to climate change is a major research objective in biogeography, evolutionary ecology and conservation (Bellard et al., 2012; Cardinale et al., 2012; Hoffmann et al., 2015) and a necessary step to design better species conservation actions and strategies. Yet, we need to factor in intraspecific genetic variation since a species is an aggregate of different genetic lineages which differ in their adaptation to environmental conditions. However, the effects of climate change on genetic diversity are just beginning to be explored (Bellard et al., 2012). Indeed, climate change can profoundly affect species distributions (Bakkenes et al., 2002; Thomas et al., 2004; Thuiller et al., 2005) as it has already been documented (Davis & Shaw, 2001; Parmesan & Yohe, 2003; Parmesan, 2006). As temperature is expected to rise over the 21^st^ century under all studied emission scenarios (IPCC, 2014), it will alter species distribution (Loarie et al., 2009). We need to predict these changes if we want to take adequate actions to mitigate future losses of biodiversity (Pfenninger et al., 2012). There are abundant examples in the scientific literature of tools and methodologies aimed at predicting species distributions in the face of global climate change (e.g. Aitken et al., 2008; Guisan & Zimmermann, 2000; Franklin et al., 2012; Pfenninger et al., 2012; Razgour 2015). However, most studies have focused on forecasting distributions at the species level and not taking into account for species internal genetic variability (Bálint et al., 2011; Benito-Garzón et al., 2011; Alsos et al., 2012; Pauls et al., 2013). Within a species, this genetic variability manifests itself as an assemblage of lineages or clusters differing in their degree of adaptation (Sork et al., 2010; Neiva et al., 2015). Such genetic clusters can be inferred from genetic markers such as microsatellites or single-nucleotide polymorphisms (SNPs) using clustering algorithms such as STRUCTURE (Pritchard et al., 2000). Hence, distribution models at the species level will be necessarily less precise than models at the intraspecific genetic level. The knowledge of this intraspecific genetic diversity may give quite a different vision on the projected distributions of species under climate change (Jump et al., 2009; Benito-Garzón et al., 2011; Yannic et al., 2014; Marcer et al., 2016) and be fundamental in conservation.

Data on intraspecific genetic lineages come as proportions (i.e. percentages of genetic cluster or lineage membership for admixed populations) instead of the presence/absence data used in traditional SDM techniques. However, despite the existence of notable studies that try to infer species distributions by working at the genetic level (e.g. Razgour et al, 2013; Valladares et al., 2014; Yannic et al, 2014), few studies (e.g. Jay et al., 2012) have directly used these proportion-type data to model the distribution of intraspecific units as suggested by Gotelli & Stanton-Geddes (2015). More studies are much in need in this respect.

A model species well suited for exploring this issue is the annual plant *Arabidopsis thaliana* (L.) Heynh *(Brassicaceae).* Indeed, for *A. thaliana*, an extensive georeferenced database has already been collected for the Iberian Peninsula (Picó et al., 2008; Manzano-Piedras et al., 2014) which is a major area of genetic diversity in the global distribution of this species (Brennan et al., 2014). The Iberian Peninsula is an environmentally diverse region where *A. thaliana* shows local adaptation to the different environments as this region was a glacial refugium (Picó et al., 2008). It comprises a wide range of climates and *A. thaliana* shows great variation in response to these different conditions (Bouchapke et al., 2008; Picó et al., 2008; Hancock et al., 2011; Fournier-Level et al., 2011; Manzano-Piedras et al., 2014). This local adaptation is reflected in the intraspecific genetic clusters of this species (Hancock et al., 2011; Fournier-Level et al., 2011).

Here, we analyse how the distribution of *Arabidopsis thaliana* could be affected by climate change at the intraspecific genetic level in the Iberian Peninsula. We used *admixed* genetic lineages inferred from a set of 250 SNPs analysed in 274 accessions spread across the Iberian Peninsula. Our objectives are *a)* to assess the effects of climate change on genetic lineages at the *admixed* population level by directly modelling genetic cluster membership values and *b*) to predict potential genetic cluster membership values across the Iberian Peninsula. Our work complements and expands comparable studies (Jay et al., 2012; Gotelli & Stanton-Geddes, 2015) and aims at modelling genetic diversity directly from values of membership to genetic clusters.

## MATERIALS AND METHODS

### Study area

The Iberian Peninsula is located in the western part of Eurasia (ca. between 10 W and 4 E, and 36 S and 44 N). It is a mountainous region of circa 580 000 km^2^ and one of the major biodiversity hotspots in the Mediterranean. It harbors around 50% of the European plant species and 31% of the European endemic plants (Williams et al., 2000; Araújo et al., 2007) and has been a major Pleistocene glacial refugium (Hewitt, 2001; Gomez and Lunt, 2006).

### Species, accessions and genetic data

We used a dataset of 274 accessions of *Arabidopsis thaliana* (L.) Heyhn. derived from the one described in Marcer et al. (2016) and which includes additional climate predictors. Genetic data consist of percentages of cluster membership assignment derived with the STRUCTURE algorithm (Pritchard et al., 2000), version v.2.2, from a set of 250 nuclear genetic polymorphisms (SNPs) as described in Manzano-Piedras et al. (2014).

## Climate data

### Present time

A set of eight bioclimatic variables relevant to the species’ ecology were selected as model predictors: BIO1 (Annual Mean Temperature), BIO2 (Mean Diurnal Range), BIO3 (Isothermality), BIO4 (Temperature seasonality), BIO8 (Mean Temperature of Wettest Quarter), BIO12 (Annual precipitation), BIO15 (Precipitation seasonality) and BIO18 (Precipitation of Warmest Quarter) (see Table ST1 in Supplementary Information). These variables were derived from the Digital Climatic Atlas of the Iberian Peninsula (http://opengis.uab.es/wms/iberia/enindex.htm) using the *dismo* package in R (Hijmans et al., 2015). Data were accessed on February 19, 2015. Their pairwise degree of collinearity (Spearman’s correlation coefficient) is lower than 0.7.

### Future scenarios of climate change

We selected the 2070 RCP2.6 and RCP8.5 climate change scenarios (Moss et al., 2008), which represent the minimum and maximum trends in radiative forcing (van Vuuren et al., 2011) of the four RCP scenarios adopted by IPCC in its fifth Assessment Report (AR5) (IPCC, 2014) as modelled by four different climate change models: HadGEM2-ES (Met Office Hadley Centre, UK), MRI-CGCM3 (Meteorological Research Institute, Japan), MIROC-ESM (Japan Agency for Marine-Earth Science and Technology, Atmosphere and Ocean Research Institute (The University of Tokyo), and National Institute for Environmental Studies), NorESM1-M (Norwegian Climate Centre, Norway). Data were downloaded from the WorldClim website (http://www.worldclim.org) on February 19, 2015. We reprojected them from their original WGS84 Lat/Long (EPSG:4326) projection into ETRS89/LAEA (EPSG: 3035) equal-area projection and resampled them from 30 seconds to 1 km resolution using bilinear interpolation. Finally, for each of the two RCP scenarios we averaged the four models to generate our GCC dataset for predictive purposes. We will refer to these simply as RCP2.6 and RCP8.5.

## Modelling approach

We used two different statistical models, namely, a parametric beta regression and a non-parametric regression tree algorithm, as implemented in R packages *betareg* and *mvpart*, respectively (Cribari-Neto & Zeileis, 2010; Therneau & Atkinson, 2014). This combination of very different techniques allowed us to check for agreements and disagreements between them to make predictions more robust. We are assuming that populations are, at least, partially adapted to their local climate conditions and that their admixture setup is not exclusively due to demographic processes; a reasonable assumption for Arabidopsis thaliana (Picó et al., 2008;

Hancock et al., 2011; Fournier-Level et al., 2011). We modelled each genetic cluster separately, using their cluster membership percentages as dependent variables and the above mentioned climate variables as independent variables. No interaction terms were used. The model formula is as follows:

Y_[GC1, GC2, GC3, GC4]_ = ß_0_ + ß_1_·BIO1 +ß_2_·BIO2 + ß_3_·BIO3 + ß_4_·BIO4 + ß_5_·BIO8 + ß_6_·BIO12 + ß_7_·BIO15 + ß_8_·BIO18

The dependent variable Y is the cluster membership coefficient for each of the four genetic clusters and can take values ranging from 0 to 1 which express the admixed degree of membership to the given genetic cluster. ß_0_ through ß_8_ are the regression coefficients and BIO1/2/3/4/8/12/15/18 are the climate predictors.

To validate models and prevent overfitting, we used 10-fold cross-validation and used the average of the mean absolute error and of the pseudo-*R*^2^ of test folds as measures of predictive performance. Finally, we fitted the models again using the whole dataset and used these final models to make predictions. For each model, we calculated Moran’s *I* at five distance intervals (2000, 4000, 6000, 8000 and 10 000 m) in order to assess residual spatial autocorrelation assessing its significant using 10000 randomizations (R package *ncf* (Bjornstad, 2013)). Spatial autocorrelation was calculated both at the variable level (vSAC) and at the models’ residuals level (rSAC) in order to evaluate whether models managed to decrease already present vSAC.

## Prediction under global climate change

### At population level

We calculated the cluster membership percentage change as predicted by each modelling method for each population and genetic cluster. We then checked for each population and climate change scenario whether both modelling methods agreed on the direction of change; *i.e.* whether a given cluster membership value either increases or decreases. For those populations in which models did not agree we did not trust the predicted changes in genetic cluster membership percentages. For the rest of populations in which the models did agree we predicted the future genetic cluster membership value as the weighted average pseudo-*R*^2^ of both modelling techniques. After that, we assessed the relative dominance of each genetic cluster in each population in order to highlight those populations for which their dominant genetic cluster changed. Finally, we assessed the degree of intraspecific composition populational change between current and future climate conditions by using the Pearson correlation coefficient between current and future genetic cluster membership percentages to give a global measure of population structure change, as in Jay et al. (2012).

### At Iberian Peninsula level

We projected both models to the whole of the Iberian Peninsula at present time climate conditions and under both RCP2.6 and RCP8.5 climate change scenarios. As with the predictions at population level we used the weighted average pseudo-*R*^2^ of both predictions to give a final map of potential genetic cluster membership suitability under climate change. We considered as unknown those areas similar to the environmental conditions of the populations for which models did not agree (greyed-out areas in Figure 3). We used the Gower’s dissimilarity coefficient (Gower, 1971) as implemented in R package *StatMatch* (D’Orazio, 2015) and determined as unpredictable those areas with a value below 0.025.

### Changes in potential distribution

In order to quantify the potentially suitable area for each genetic cluster and to compare its extent between present time and climate change scenarios, we used a threshold of 0.5 as a cut-off point for determining whether any given location is either suitable or unsuitable for each genetic cluster. We then calculated the potentially suitable area for each genetic cluster and scenario of climate change as well as additional measures of change such as the mean and median of the suitability scores.

## RESULTS

### Model performance and residual spatial autocorrelation

Overall, regression trees models showed consistent better predictive performance than beta regression ones for both mean absolute error (mae) and pseudo-*R*^2^ (hereafter pr^2^) metrics (Table ST2). For all genetic clusters, beta regression results in an average mae of 0.158 and average pr^2^ of 0.263 while regression trees result in an average mae of 0.126 and average pr^2^ of 0.320. The best predicted genetic cluster was GC2 for both regression trees (mae: 0.081 ± 0.013, pr^2^: 0.561 ± 0.142) and beta regression (mae: 0.103 ± 0.016, pr^2^: 0.468 ± 0.142). On the other hand, GC3 had the highest error in the case of beta regression models (mae: 0.205 ± 0.027, pr^2^: 0.103 ± 102), while for regression trees it was GC1 according to mae (0.163 ± 0.025) and GC3 according to pr^2^ (0.144 ± 0.116).

Tables ST3-1/4 and figures SF3-1/4 show results of the spatial autocorrelation (SAC) analysis. At the variable level (vSAC), it was always highest at the second distance class (4 000 m) for all genetic clusters with the exception of GC4, for which it was the third distance class (6 000 m). Except for 5 cases (GC1-Beta-4000, GC2-Beta-4000, GC2-RT-4000, GC4-Beta-8000 and GC4-RT-8000) residual SAC (rSAC) was always lower than vSAC, meaning that the models were able to reduce inherent vSAC in most cases. For GC1 and GC4 rSAC was not significant in all models and distance classes except for one case each. GC3 rSAC was significant in two distance classes of the beta model and one distance class of the regression tree model. Beta regression for GC4 only had one class with non-significant rSAC but the regression tree model rSAC was only significant in two of them.

### Predictions at the population level

#### Model comparison and prediction of change for genetic cluster membership percentages

Table 1 and Figure 1 summarize the degree of agreement between both modelling methods in determining the direction of change in genetic cluster membership percentages per population and climate change scenario. In most of the populations and for both climate change scenarios, both beta regression and regression tree models coincide in determining the direction of change, i.e. whether a given population increases or decreases its membership percentage for a given genetic cluster. For scenarios RCP2.6 and RCP8.5 the maximum coincidence was in the case of GC1, for which they only differed in 25 populations (9.1 %), while the minimum coincidence for RCP2.6 was in the case of both GC2 and GC4 with 50 populations each (18.2 %), and for RCP8.5 was in the case of GC4 with 63 populations (23.4 %). Considering only the populations for which models agreed in the direction of change, we can state that for scenario RCP2.6, GC1 had the biggest gain (+0.169 ± 0.100) and GC3 the biggest loss (-0.456 ± 0.273), while for scenario RCP8.5 the biggest gain was for GC3 (+0.214 ± 0.172) and the biggest loss also for GC3 (-0.434 ± 0.268). In figure 1 populations for which models do not agree are marked with a black diamond.

**FIGURE 1.**
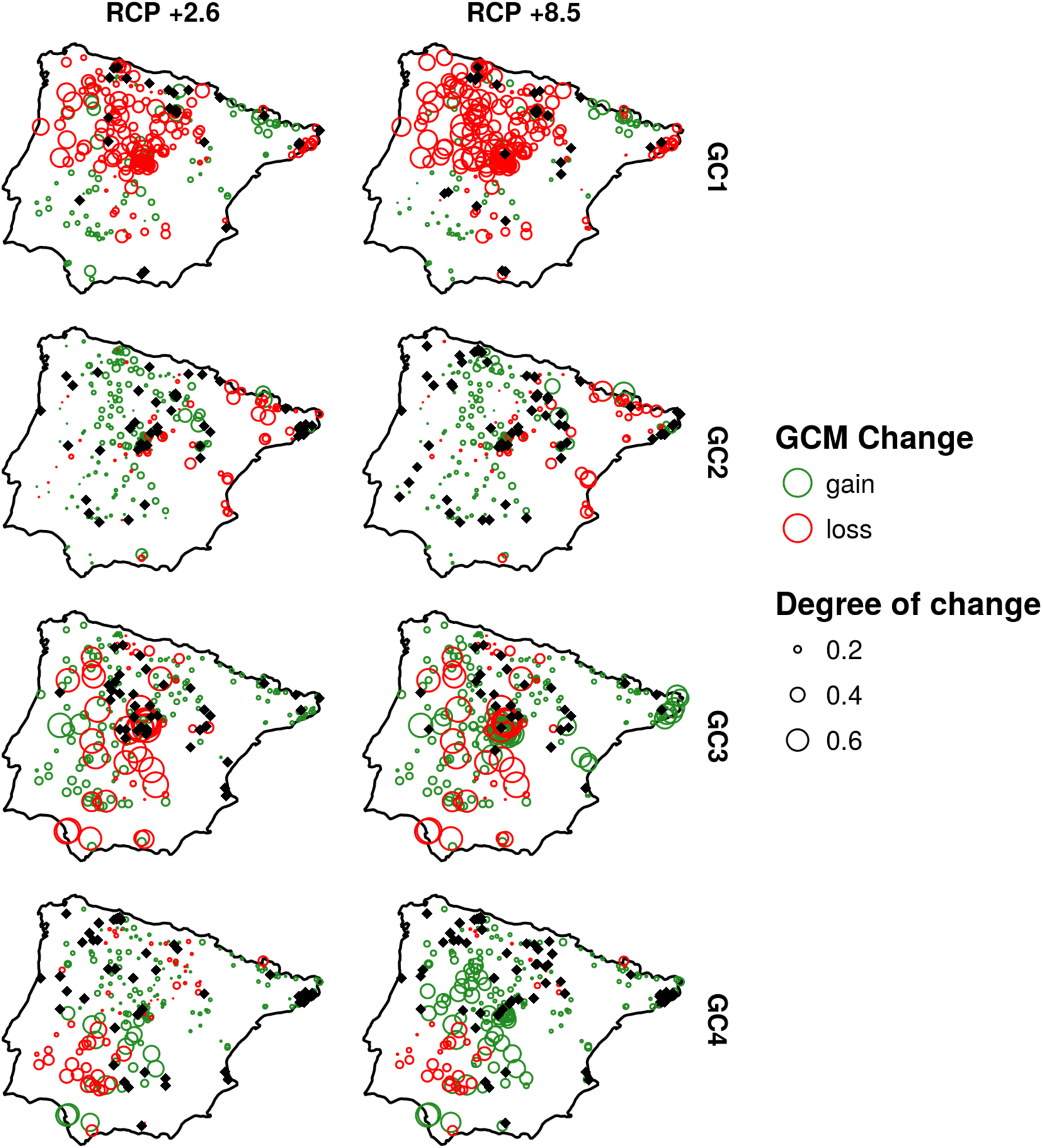
Gains and losses in membership percentage to genetic clusters. Black diamonds show the populations for which models do not agree. Degree of change is expressed in membership percentage units divided by 100.

**TABLE 1.** Changes in genetic cluster membership at the population level. Columns - gen.unit: Genetic cluster, rcp: Representative concentration pathway (climate change scenario), n.gain: Number of populations for which models agree on an increase in genetic cluster membership value, n.loss: Number of populations for which models agree on a decrease in genetic cluster membership value, unknown: Number of populations for which models do not agree, avg.gain: Average gain for populations on which models agree, sd.gain: Standard deviation of gain for populations on which models agree, avg.loss: Average loss for populations on which models agree, sd.loss: Standard deviation of loss for populations on which models agree

| gen.unit | rcp | n.gain | n.loss | unknown | avg.gain | sd.gain | avg.loss | sd.loss |
| --- | --- | --- | --- | --- | --- | --- | --- | --- |
| GC1 | +2.6 | 93 | 156 | 25 | 0.169 | 0.100 | -0.264 | 0.127 |
| GC1 | +8.5 | 76 | 173 | 25 | 0.138 | 0.084 | -0.335 | 0.160 |
| GC2 | +2.6 | 157 | 67 | 50 | 0.117 | 0.078 | -0.136 | 0.098 |
| GC2 | +8.5 | 141 | 72 | 61 | 0.122 | 0.089 | -0.148 | 0.109 |
| GC3 | +2.6 | 183 | 50 | 41 | 0.135 | 0.074 | -0.456 | 0.273 |
| GC3 | +8.5 | 200 | 46 | 28 | 0.214 | 0.172 | -0.434 | 0.268 |
| GC4 | +2.6 | 161 | 63 | 50 | 0.129 | 0.110 | -0.162 | 0.124 |
| GC4 | +8.5 | 173 | 38 | 63 | 0.200 | 0.138 | -0.190 | 0.107 |

### Genetic cluster turnover

Figure 2 shows for each climate change scenario and genetic cluster which populations are predicted to have their dominant genetic cluster changed. Numbers are calculated taking only into account those populations for which models agree (black dots in Figure 2), which is not applicable to present time. All genetic clusters but GC4 will suffer a switch in suitability for some of their dominant populations; *i.e.* their present time dominant cluster will be switched to another one. Overall, taking into account all shifts from every genetic cluster at present time, GC1 will go from 148 dominated populations to 132 in RCP2.6 and 60 in RCP8.5 with 16 populations of uncertain fate in RCP2.6 and 12 in RCP8.5. GC2 will go from its current 57 populations to 43 in RCP2.6 and 58 in RCP8.5 with 18 populations of uncertain fate in RCP2.6 and 15 in RCP8.5. GC3 will go from its current 38 populations to only 9 in RCP2.6 and 55 in RCP8.5 with no populations of uncertain fate in RCP2.6 and 3 in RCP8.5. Finally, GC4 will be the only genetic cluster which will keep all its present time dominated populations in both climate change scenarios and will increase from its current 31 to 51 in RCP2.6 (with 5 of uncertain fate) and to 67 in RCP8.5 (with 4 of uncertain fate).

**FIGURE 2.**
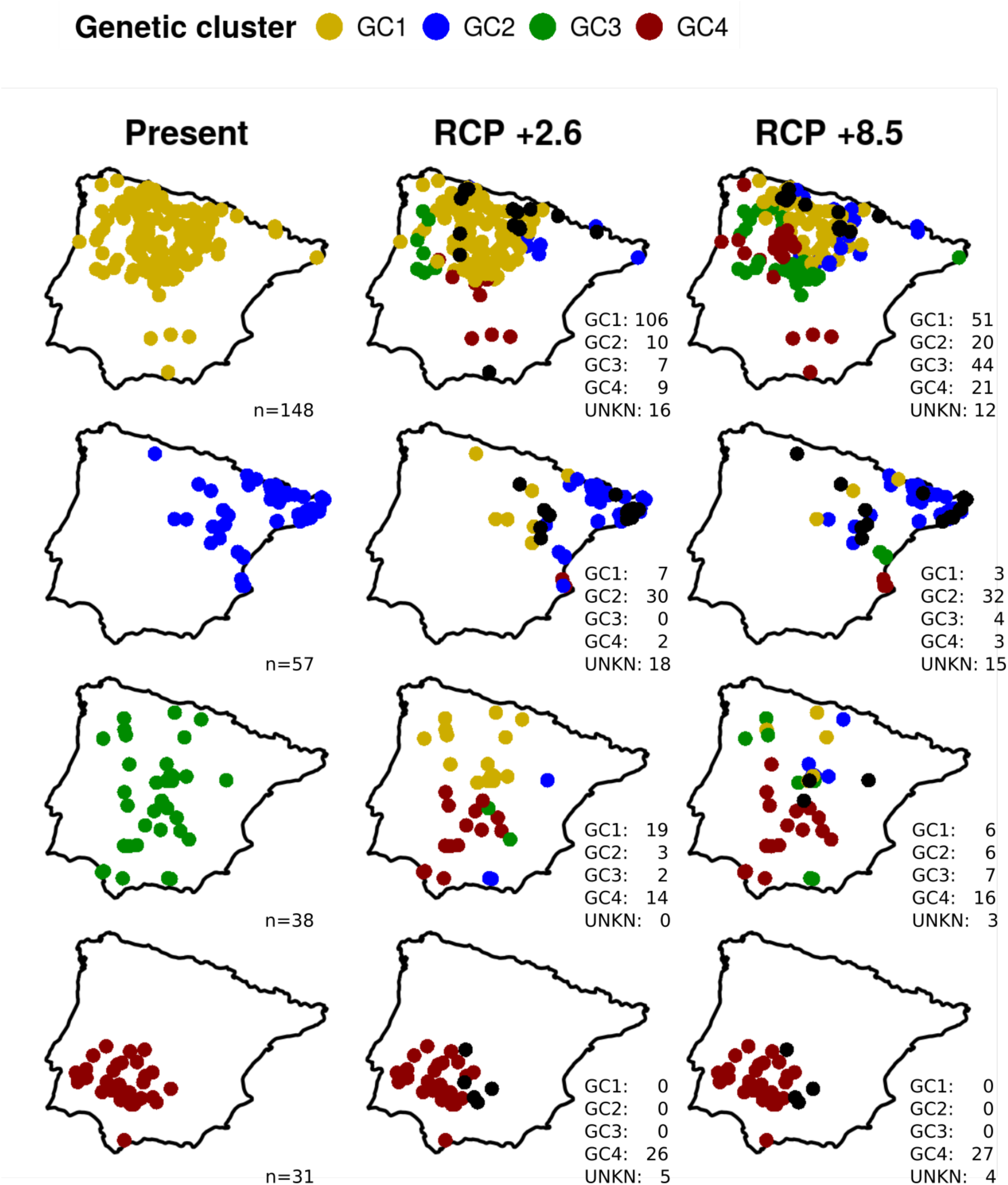
Genetic cluster turnover for populations under two scenarios of climate change in 2070. Numbers at bottom right of each map show the number of populations dominated per genetic cluster in each case. UNKN (unknown) is the number of populations for which models do not agree.

Figure SF4 and SF5 show the degree of genetic structure change that each population must undergo to be best suited for its new environment. Approximately a fifth of existing populations will have to undergo a major structural change: 17.3% in the case of RCP2.6 and 21% in the case of RCP8.5 (Table ST4).

## Predictions at the level of the Iberian Peninsula

Table 2 shows the potentially suitable area, the mean and the median for each genetic cluster at present time and future scenario as a way to measure the overall change across the Iberian Peninsula (Figure 3) that each genetic cluster would undergo in each scenario of climate change. Genetic clusters GC1 and GC2 show a reduction in mean suitability. For RCP2.6 their means get reduced from 0.314 to 0.270 and from 0.208 to 0.204 and their potential distribution area gets reduced by 50.8% and 87.5%, respectively. On the other hand, GC3 and GC4 would undergo a general increase in their suitability to climate change. Although GC3 would decrease its mean from 0.225 to 0.217 for RCP2.6, for RCP8.5 it would increase up to 0.257. With respect to potential distribution area, it would increase by 835.0% for RCP2.6 and by 4887.4% for RCP8.5. Finally, land suitability for GC4 would also be substantially increased; from a mean value of 0.253 to 0.302 for RCP2.6 and up to 0.369 for RCP8.5 and an increase of potential distribution area of 175.2% for RCP2.6 and of 238.1% for RCP8.5.

**FIGURE 3.**
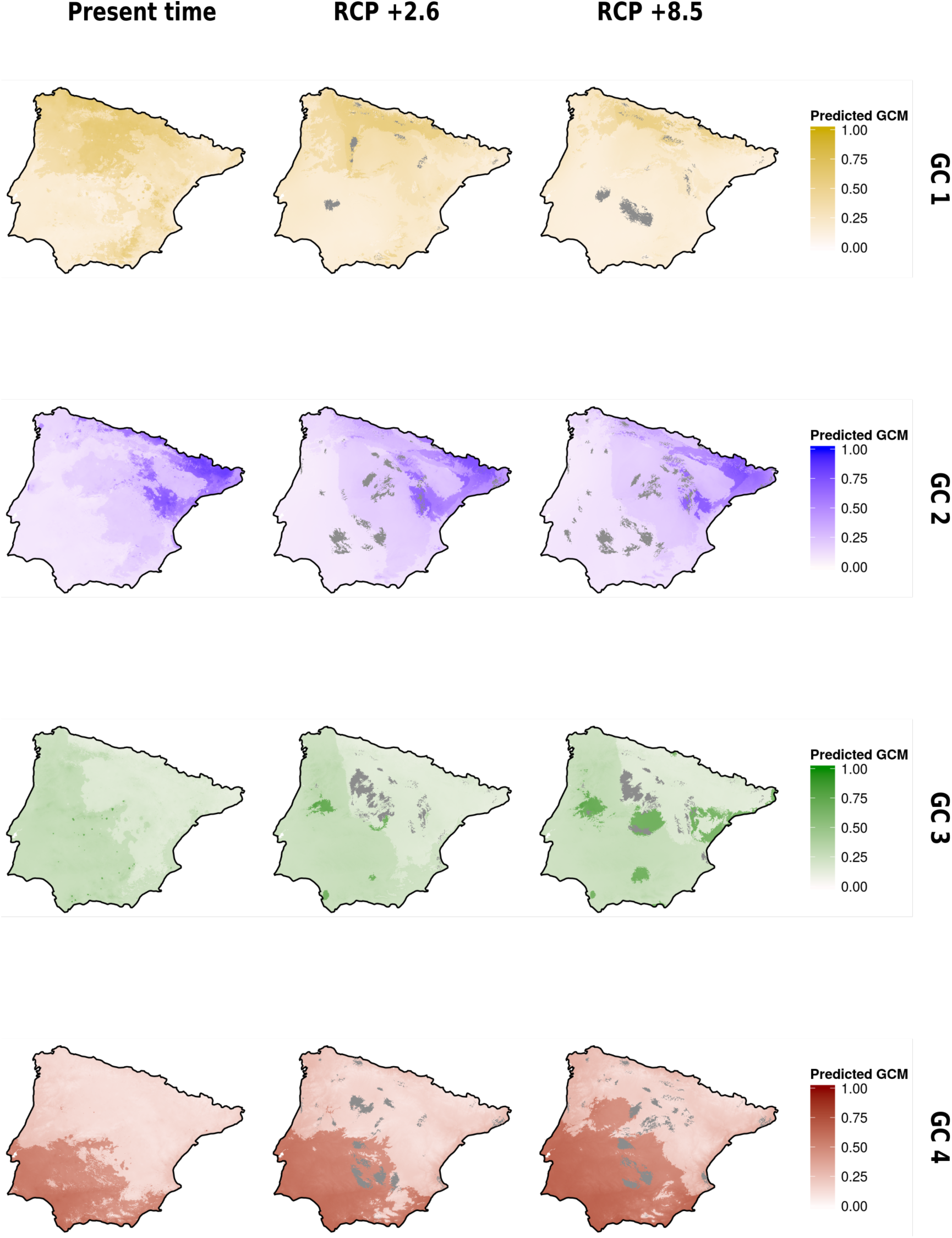
Projection of genetic cluster membership percentage under two scenarios of climate change (2070). Gray areas are zones environmentally similar to the environmental conditions of the populations for which models do not agree and thus cannot be predicted with confidence. Legends are expressed in membership percentage values divided by 100.

**TABLE 2.** Changes in land suitability in the Iberian Peninsula for each genetic cluster and time frame. Potential area is expressed in km^2^ after applying a cut-off threshold of 0.5. Mean and Median are the mean and median predicted cluster membership percentage divided by 100, respectively. Present time shows the current values for each Genetic cluster and Statistic, and RCP +2.6 and RCP +8.5 show these values for each climate change scenario.

| Genetic cluster | Statistic | Present time | RCP +2.6 | RCP +8.5 |
| --- | --- | --- | --- | --- |
| GC1 | Potential area | 101900 | 50173 | 12713 |
| GC1 | Mean | 0.314 +/- 0.148 | 0.270 +/- 0.113 | 0.194 +/- 0.088 |
| GC1 | Median | 0.290 | 0.224 | 0.162 |
| GC2 | Potential area | 60870 | 48371 | 42885 |
| GC2 | Mean | 0.208 +/- 0.167 | 0.204 +/- 0.154 | 0.198 +/- 0.146 |
| GC2 | Median | 0.162 | 0.178 | 0.171 |
| GC3 | Potential area | 1033 | 8630 | 50487 |
| GC3 | Mean | 0.225 +/- 0.062 | 0.217 +/- 0.080 | 0.257 +/- 0.132 |
| GC3 | Median | 0.251 | 0.231 | 0.244 |
| GC4 | Potential area | 89036 | 156021 | 212014 |
| GC4 | Mean | 0.253 +/- 0.168 | 0.302 +/- 0.180 | 0.369 +/- 0.188 |
| GC4 | Median | 0.154 | 0.200 | 0.300 |

The spatial distribution of these changes is shown in Figure 3 where there are clear trends northward. Higher values of suitability for GC1 get more constrained to the most northern part of the Iberian Peninsula both for RCP2.6 and RCP8.5. GC2 shows a shift towards the north-east of its climate suitability. GC3 moves towards central parts of the Iberian Peninsula and GC4 again shows a clear northerly expansion of climate suitability. Supplementary materials provide Figures SF6 and SF7-1/4 which show future predictions differentiated by modelling method.

## DISCUSSION

Our results confirm the need for dealing with intraspecific genetic variation in order to understand and forecast the effects of climate change on species distributions as has been suggested (Hancock et al., 2011; Fournier-Level et al., 2011). We have shown that the impact of climate change is unevenly felt by the different genetic lineages of *Arabidopsis thaliana* in the Iberian Peninsula, an expected result given that these lineages are known to have local adaptation and their relative percentage mixture in populations is geographically structured in the Iberian Peninsula (Picó et al., 2008). Moreover, it is possible to identify those populations which, for their particular genetic makeup, will need to undergo a major structural genetic change from those which may have a minor impact. This is a potentially useful outcome since it can help in optimizing scarce resources when undertaking conservation efforts.

Species distribution models should therefore consider genetic variation as has been suggested (Jump et al., 2009; Benito-Garzón et al., 2011; Yannic et al., 2016; Marcer et al., 2016). Although for the vast majority of species this kind of information is unavailable and traditional SDM techniques might be the only option, these results clearly suggest that, when possible, this should be the way to proceed. Also, given the fast pace in which genetics and genomics technologies progress (Hoffman et al., 2015; Tyler-Smith et al., 2015) it is not unthinkable that in a not so distant future this could be done for most species.

It is important to recognize that these kinds of analyses forget an important component, demography. As shown, we can give a snapshot of what might happen to a given species in the face of rising temperatures and shifting climate regimes. However, this is done as a single leap forward into the future without considering the population dynamics and resulting demographic changes which would occur if a sequence of many events in a real-time path to the projected 2070 year was considered. Although future research should try to incorporate demographic processes at the landscape level into the modelling process, this is not an easy task. The difficulty resides not only in model development but more importantly in the lack of sufficient quality data (Ehrlén & Morris, 2015) on species life history traits and populations and in the stochasticity, inherent to some processes like long-distance dispersal events (Pergl et al., 2011). In order to project spatially explicit models in time we need data on local demographic parameters over broad areas (Nathan 2006; Thuiller et al., 2014), which is usually unavailable. Even in the case of *Arabidopsis thaliana*, a model species and one of the better-known species, there is no reliable field-obtained data on important traits, such as population size, seed production, seedling survival or dispersal distance of natural populations over broad areas, which would be needed to build such models. A possibility is that of running simulations but this has it’s owns caveats, too. Without actual data, their outcomes are very dependent on the value of the parameters that are fed to them and, thus, their results difficult to interpret.

The expected effect of climate change on the genetic lineages of *Arabidopsis thaliana* in the Iberian Peninsula is in accordance to the work of Marcer et al. (2016). Genetic clusters GC3 and GC4 will increase their potential distribution by expanding northward their distribution ranges. On the other hand, GC1 and GC2 will shrink their distribution ranges and their highest suitable areas will be more constrained to the north and north-east, respectively. At the population level, *Arabidopsis thaliana* will undergo a genetic turnover in many of its populations which will need to shift from one dominant genetic cluster to another. Either dispersal, local re-adaptation or phenotypic plasticity must come to the rescue if most populations are to survive these changes induced by climate change. For instance, as shown by Picó et al. (2014), flowering time, which is mediated by temperature, appears to be the major life-history trait adjusting *Arabidopsis thaliana* to the different environmental conditions across the Iberian Peninsula. Plasticity in this and other traits may provide for some necessary time for populations to adapt (Donohue et al., 2005; Levin, 2009; Nicotra et al., 2010).

We have modelled genetic cluster membership percentages to infer potential future distributions at the intraspecific level rather than resorting to the use of thresholds to transform data into presence/absence and use SDM techniques. This work adds to the need for more research in this respect (Jay et al., 2012; Gotelli & Stanton-Geddes, 2015). We used two very different modelling approaches, a parametric beta regression and a non-parametric regression tree analysis, which showed a high degree of agreement in their predictions. Despite, their individual predictive power with a set of only-climate predictors was limited, their combined use allowed us to identify which predicted changes could be trusted and use this information to make predictions with highlighted populations and areas of uncertainty. The limited predictive power should not be attributed to the statistical modelling techniques themselves but it probably reflects the lack of important predictors such as land use and soil type (Marcer et al., 2016), a price currently needed to pay due to the lack of climate change models for these types of variables.

Finally, we would like to express the need for quality data if we are to understand the biological and environmental processes that drive species distributions and predict the fate of species in front of climate change. The building of quality and extensive datasets on natural populations such as the one used in this study require substantial resources in terms of time and funding. Yet, they are paramount to such undertakings. The lack of sufficient funding for such basic research and of community recognition for data building and curation is a major handicap which hinders advancement in this respect. The authors encourage funding agencies not to dismiss this pressing need.

## ACKNOWLEDGEMENTS

A.M. and M.J.F were partially supported by the NEWFORESTS project (PIRSES-GA-2013-612645) from the European 7FP. F.X.P. and A.M. were partially supported by Ministerio de Economía y Competitividad of Spain (CGL2012-33220/) and the Agency for Management of University and Research Grants of the Generalitat de Catalunya (2014-SGR-913). F.X.P. acknowledges the Severo Ochoa Programme for Centres of Excellence in R+D+I (SEV-2012-0262) in Spain.

